# ARID1A deficiency unleashes centromeric RNA transcription to drive chromosomal instability and boosts PKMYT1 inhibitor efficacy via RNA sensing

**DOI:** 10.64898/2026.04.11.717899

**Authors:** Chengguo Li, Xueqian Cheng, Weizhen Liu, Gengyi Zou, Thi Hong Minh Nguyen, Emily Zhao, Noemi Ira, Liyong Zeng, Yibo Fan, Shilpa Dhar, Guocan Wang, Shumei Song, Ming Zhao, Rugang Zhang, Yuan-Hung Lo, Timothy A. Yap, Jaffer Ajani, Guang Peng

## Abstract

Cancer gene-associated mutations and molecular hallmarks of chromosomal instability (CIN) are unexpectedly common in histologically normal cells and tissues. These emerging findings challenge the binary distinction between “normal” and “cancerous” cells and suggest that early tumorigenesis may commence against a background of widespread yet largely tolerated genomic instability. However, it remains largely unexplored how a cancer gene-associated mutation can initiate the development of CIN-like states in non-malignant cells and drive tumor evolution. *ARID1A*, a chromatin remodeling factor, was identified as the most frequently mutated gene in both gastric normal epithelium and tumors. This distinctive molecular convergence presents an opportunity to elucidate the mechanisms by which a cancer-associated gene facilitates the initiation of early CIN phenotypes and develop effective antitumor strategies. In the present study, using primary human gastric organoids, we employed optical genome mapping (OGM) and live-imaging technologies to demonstrate that *ARID1A* depletion induced a wide spectrum of structural variants (SVs), copy number variants (CNVs), and chromosomal segregation errors, characteristic features of CIN at a very early stage of gastric tumorigenesis. Mechanistically, ARID1A bound centromere repetitive satellite DNA (satDNA) sequences. Its SWI/SNF-associated chromatin remodeling activity was required for suppressing satDNA transcription and the production of α-SatRNA, through restricting RNAPII elongation. Consequently, *ARID1A* depletion led to overexpression of α-SatRNA, and a higher incidence of sister chromatid exchange (SCE), a sensitive indicator of CIN. Importantly, the elevated α-SatRNA expression in *ARID1A*-deficient cells further established a dual therapeutic vulnerability for G2/M checkpoint blockade, such as PKMYT1 inhibitor (PKMYTi), by concurrently aggregating CIN-induced cell death and activating self-dsRNA sensing-mediated innate immune response. Notably, PKMYTi markedly promoted α-SatRNA expression, aberrant release of these self-derived dsRNAs into the cytosol and a robust activation of the RIG/MDA5-MAVS-depenent type-I interferon response in *ARID1A*-depleted cells. As expected, PKMYTi potentiated the efficacy of immunotherapy in *ARID1A*-deficient gastric tumors. Together, our findings reveal that *ARID1A* deficiency unleashes centromeric α-SatRNA transcription, which sets the molecular stage for tumor evolution and targeted therapy by coordinately inducing CIN and self-dsRNA-induced innate immune responses.

## Introduction

It is well established that somatic mutations of cancer-driver genes are prevalent in “normal” tissues ^1–8^. What has been less clear is how these mutations identified in normal tissues set the stage for cancer to evolve. A question of more clinical significance is how these mutations affect cellular functions to prime and shape the cancer hallmarks, and whether this understanding could inform strategies for cancer prevention and treatment.

In addition to somatic mutations, a most recent study of normal gastric epithelium reported that single-nucleotide variants (SNVs), somatic structural variants (SVs) and copy number variants (CNVs), key features of chromosomal instability (CIN), are found in normal gastric glands at considerably higher prevalence than in other normal human cell types reported so far^2,4–6^. Very unusually for normal cells, gastric epithelial cells frequently exhibit recurrent trisomies of specific chromosomes, another hallmark of CIN primarily attributed to chromosome segregation abnormalities^6^. More strikingly, the persistence of these large-scale genomic alterations of CIN is estimated from a very early age around adolescence, which is much earlier than the accumulation of somatic mutations linearly through normal tissue ageing^6^. Furthermore, although the presence of CIN is not significantly linearly associated with age, it is significantly enriched in the context of severe chronic inflammation^6^. CIN has long been recognized as a hallmark of cancer^9–11^. Evidence strongly links CIN to tumor evolution and tumor-promoting inflammation^12^. However, the molecular changes that induce CIN and its associated inflammation in such an early stage of gastric cancer evolution remain unknown.

*ARID1A* (the AT-rich interaction domain 1A), also known as BAF250a, is a subunit of the chromatin remodeling complex SWI/SNF (mating type SWItch/Sucrose NonFermentable). This complex uses ATP hydrolysis to regulate chromatin structure and DNA accessibility to proteins involved in chromatin transactions, such as gene transcription, DNA repair, DNA replication, and telomere maintenance^13^. *ARID1A* is widely recognized as one of the most extensively mutated genes in cancers^14,15^. The majority of these mutations are inactivating in nature, resulting in the loss of *ARID1A* expression^14^ ^15^. Following *TP53*, *ARID1A* is the second most mutated gene in gastric cancer (GC), exhibiting a mutation rate of approximately 30%^16,17^. Most intriguingly, *ARID1A* is identified as the most frequent somatic ‘driver’ mutation of known cancer genes in normal gastric epithelium^6^. These findings raise critical questions regarding whether and how somatic mutations of cancer genes in normal cells, such as *ARID1A*, may induce CIN and trigger inflammation signaling, the two fundamental hallmarks of cancer that set the stage for tumor development. Furthermore, it is unclear whether this understanding can aid in the development of strategies to selectively target ARID1A deficiency based on the concept of synthetic lethality.

Here, we report that chromatin remodeler ARID1A is at the crossroads of regulating centromeric RNA transcription, controlling chromosome stability, and modulating innate immune response triggered by self-RNA sensing. We found that ARID1A deficiency promotes double-stranded RNA (dsRNA) transcription from centromeric regions of repetitive satellite DNAs (satRNA). These self-derived dsRNAs serve as potent molecular inducers of CIN at an early stage of tumorigenesis, and furthermore exhibit remarkable immune modulatory capabilities, effectively sensitizing tumors to the synergistic combination of G2/M cell cycle checkpoint and immune checkpoint blockades.

## Results

### ARID1A deficiency induces chromosomal instability and chromosome segregation errors in primary human gastric organoids

To study the early genetic alterations induced by ARID1A deficiency, we used a well-characterized forward-genetic human *ARID1A*-deficient organoid model, which represents a robust *in vitro* culture model to recapitulate essential attributes of primary human tissue ^18^. In this model, *ARID1A* KO was not compatible with organoid survival. Thus, *TP53* was depleted to establish an isogenic pair of *ARID1A*-wildtype (WT) (*ARID1A*^WT^/*TP53*^KO^) and *ARID1A*-KO (*ARID1A*^KO^/*TP53*^KO^) premalignant organoid lines from normal human gastric corpus using CRISPR/Cas9 technology^18^. Studies from our group and others have shown that gastric tumors with *ARID1A* mutations exhibit DNA mismatch repair deficiency (dMMR) and molecular features of microsatellite instability (MSI)^19–22^. Therefore, we sought to determine whether ARID1A loss has a molecular impact on MSI in the premalignant organoid model. Immunohistochemistry (IHC) analysis showed no significant changes in the expression of core MMR proteins including MLH1, MSH2, MSH6, and PMS2 in *ARID1A-*KO organoids compared to *ARID1A*-WT organoids (**Extended Data Fig. 1a**). Whole-exome sequencing (WES) was performed to further confirm this observation. *ARID1A-*KO organoids showed comparable levels of somatic single-nucleotide variants (SNVs) and indels as *ARID1A*-WT organoids (**Extended Data Fig. 1b**). These data suggest that *ARID1A* depletion does not alter the mutational landscape or drive MSI at the premalignant stage. This raises the question of whether *ARID1A* deficiency may induce chromosomal instability (CIN), another major form of genomic instability defined by chromosome rearrangements including structural variants (SVs) and copy number variants (CNVs) instead of SNVs. To this end, we employed optical genome mapping (OGM) technology to construct unbiased genome-wide SV and CNV maps induced by ARID1A deficiency in pre-malignant organoid models. Using the Genome Reference Consortium GRCh38/hg38 as the reference, Circos plot analysis illustrated increased SVs induced by *ARID1A* depletion including insertions, deletions, inversions, duplications, intra-chromosomal translocations, and absence of heterozygosity/loss of heterozygosity (AOH/LOH) and CNVs (**Fig. 1a**). In addition to hg38 reference genome, we further used *ARID1A*-wildtype (WT) (*ARID1A*^WT^/*TP53*^KO^) cells as a genomic baseline to subtract the potential impact of p53 loss on genomic alterations. *ARID1A*-depleted organoids (*ARID1A*^KO^/*TP53*^KO^) exhibited elevated frequencies of various chromosomal aberrations, with the most significant increases observed in AOH/LOH and CNV gain segment alterations (**Fig. 1b**). Precise mapping of CNV gains and AOH/LOH in the genomic regions is shown in **Fig. 1c**. Interestingly, clinically significant translocations t (2:8) and t (8:21), frequently implicated in lymphoma and leukemia, were also identified in *ARID1A*-depleted organoids (**Fig. 1d**). Furthermore, SV alterations were also identified in known cancer genes such as ERBB4, EGFR, ITGB8 and BRD2 (**Fig. 1e**). Notably, we did not observe whole-chromosome aneuploidy events in *ARID1A*^WT^/*TP53*^KO^ or *ARID1A*^KO^/*TP53*^KO^ organoids, indicating that whole-chromosome aneuploidy might be a late genetic event or that additional mutations are necessary for cells with whole-chromosome aneuploidy to survive. Collectively, comprehensive genome-wide SVs/CNVs profiling through OGM reveals that ARID1A deficiency induces molecular features of CIN rather than MSI in premalignant-stage organoids.

**Fig 1.**
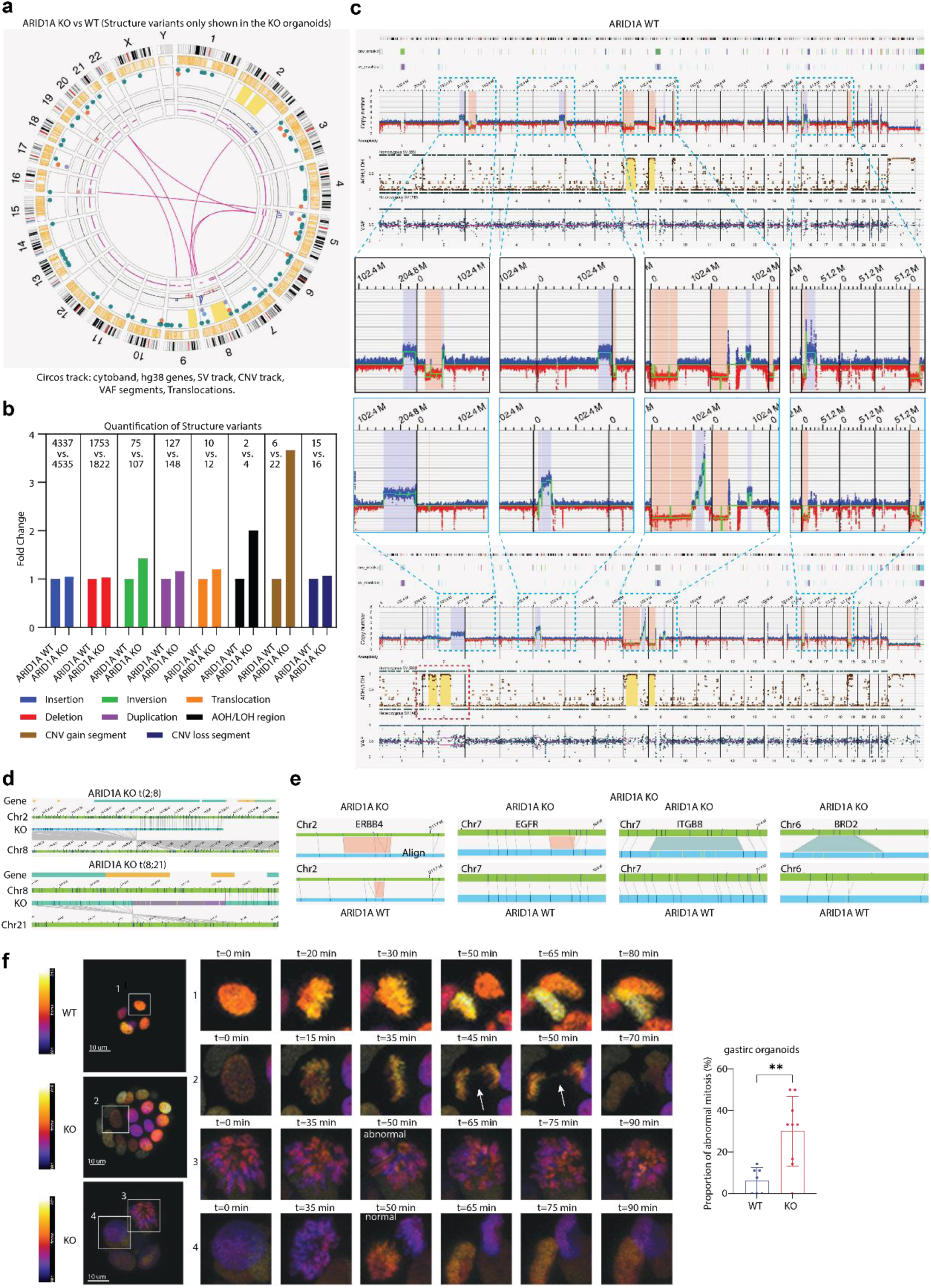
ARID1A deficiency induce chromosomal instability and mitotic errors. **a,** Circos plot of the unique structural variants detected by OGM in *ARID1A*^KO^ gastric organoid, compared to *ARID1A*^WT^ organoids. **b,** Quantification of structural variants and copy number variation (CNV) in *ARID1A*^WT^ and *ARID1A*^KO^ gastric organoids. **c,** Whole genome view of the CNV and Absence of Heterozygosity/Loss of Heterozygosity (AOH/LOH) changes in *ARID1A*^WT^ and *ARID1A*^KO^ gastric organoids. **d,** Representative translocations identified in *ARID1A*^KO^ gastric organoid. The upper panel shows chr8:161,769,408; chr2:114,898,643. The lower panel shows chr8:129,771,076; chr21:9,021,870. **e,** Representative deletions or insertions identified in *ARID1A*^KO^ gastric organoid. Deletion in ERBB4 and EGFR, and insertion in ITGB8 and BRD2 are shown. The green line is the GRCh38 genome reference. **f,** Representative images and quantification of the abnormal mitotic process in ARID1A^WT^ (n=7) and ARID1A^KO^ (n=9) gastric organoids. Student’s T test, ** P<0.01.

Given that CIN often arises from errors during cell division, we conducted live-cell imaging to investigate whether ARID1A loss impairs chromosome segregation fidelity. *ARID1A* WT and KO organoids stably expressing mCherry-tagged histone H2B (to visualize chromatin) were imaged in three dimensions at 5-minute intervals for 16-18 hours. Mitotic events were classified as either correct (no error) or in the case of obvious errors at the mid-anaphase stage as abnormal mitosis (**Fig. 1f**). Various classical mitotic errors were observed in *ARID1A*-depleted organoid cells including anaphase chromatin bridges (chromatin stretches that span the two separating packs of chromosomes and that can subsequently undergo chromosomal breakage), multipolar spindle formation (that can fragment into multiple poles), and binucleation (that can result from failed cytokinesis). In the molecular context of p53 depletion, the average percentage of abnormal mitosis was relatively low in wild-type organoids (average 6.31%), which serves as the background. In contrast, organoids with *ARID1A*-KO exhibited a significantly elevated mitotic abnormality rate of 30.11% (**Fig. 1f**). Consistent with live-imaging observations, RNA profiling data of the organoids showed that genes upregulated by ARID1A depletion were significantly enriched in chromosome segregation and mitotic pathways by gene ontology (GO) enrichment analysis (**Fig. 2a**). Gene Set Enrichment Analysis (GSEA) further confirmed that pathways involved in DNA repair, mitotic metaphase and anaphase, sister chromatid separation, the G2/M checkpoint, and mitotic spindle checkpoint were activated in *ARID1A* KO cells (**Fig. 2a**). These data suggest that *ARID1A* deficiency induces chromosome segregation errors and characteristic features of CIN in premalignant gastric organoid models.

**Fig 2.**
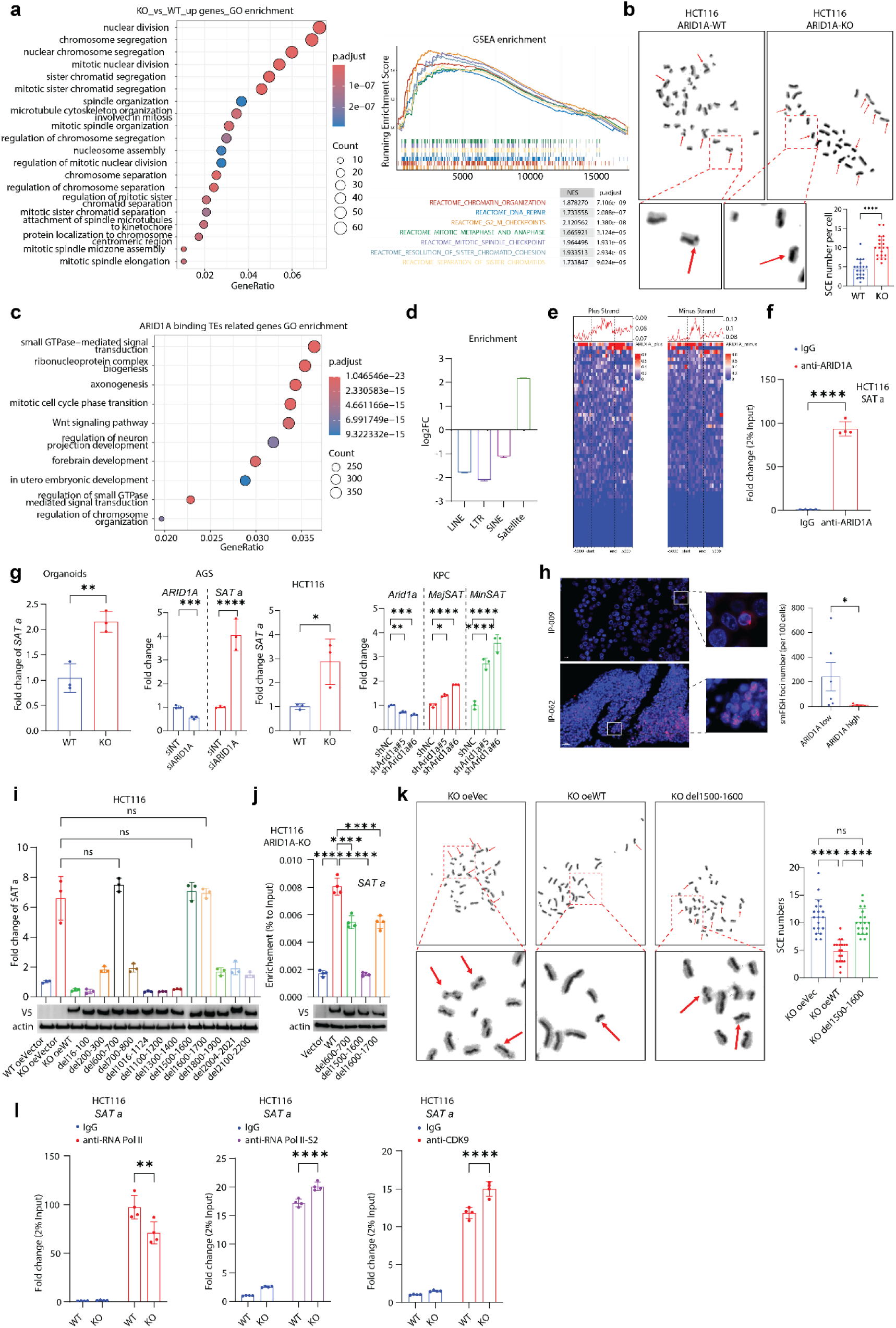
ARID1A binds to satellite alpha (α-Sat) DNA and regulates α-SatRNA expression. **a,** GO (left) and GSEA (right) enrichment results from RNA seq data of gastric organoid. For GO enrichment, the up-regulated genes (log2FoldChange>1 and FDR<0.05) verified by RNA seq was used. For GSEA, the REACTOME database and the deferentially expressed genes between *ARID1A*^KO^ and *ARID1A*^WT^ gastric organoids were used, and the top 7 pathways were shown. **b,** Representative images and quantification of sister chromatid exchange (SCE) in HCT116 *ARID1A* WT and *ARID1A* KO cells. **c,** GO enrichment of the ARID1A binding repeat sequences overlapped genes, using the dataset GSE254865. **d,** Bar plot showing the enrichment of the different types of repeat sequences that ARID1A bound. ARID1A binding peaks were re-annotated with RepeatMasker and the proportion of each type of repeat sequence was quantified (GSE254865). The enrichment was calculated as compared to the reference genome background. **e,** Heatmap showing ARID1A binding peaks to the centromere satellite DNA of the sense (ARID1A_plus) and anti-sense (ARID1A_minus) strands (GSE254865). **f,** ChIP-qPCR analysis of ARID1A protein binding to α-Sat DNA. Foldchange was normalized to the IgG (n=4). Student’s T test was used. **g,** qPCR results of α-SatRNA or MajSAT, MinSAT changes in gastric organoids (left), AGS, HCT116 (middle), and KPC (right) cells with *ARID1A*/*Arid1a* knockdown or knockout. Student’s t test or one way ANOVA was used. **h,** Representative images and quantification of smFISH staining for α-SatRNA in human peritoneal metastatic gastric cancer cells derived from ascites. ARID1A high (n=5) or low (n=6) was verified by single-cell sequencing data retrieved from EGAS00001004443. Student’s t test was used for the analysis. **i,** qPCR results of the α-SatRNA changes in HCT116 WT or *ARID1A*^KO^ cells transduced with *ARID1A* WT or indicated truncated plasmids for 72h (n=3). One-way ANOVA was used. **j,** V5-tagged ARID1A WT or indicated truncated plasmids were transduced into HCT116 *ARID1A*^KO^ cells for 72h. ChIP-qPCR was conducted with V5-antibody conjugated beads. The enrichment was calculated as proportion to the input. One-way ANOVA was used. **k,** Representative images and quantification of SCE in HCT116 *ARID1A* KO cells transduced with empty vector (oeVec), *ARID1A*-WT (oeWT), or *ARID1A*-del100-1600 (del1500-1600) plasmids for 72h (n=19). One-way ANOVA was used. **l,** ChIP-qPCR showing the enrichment of RNA Polymerase II (RNA Pol II) (left), S2P (middle), or CDK9 (right) on α-Sat DNA in HCT116 *ARID1A*-WT or KO cells. Fold change was normalized to IgG (n=4). Student’s t test was used. *P<0.05, **P<0.01, ***P<0.001, ****P<0.0001.

### The CIN phenotype induced by ARID1A deficiency is preserved in fully transformed cancer cells

We then analyzed the stomach adenocarcinoma (STAD) cohort from The Cancer Genome Atlas (TCGA) to determine whether *ARID1A* deficiency is associated with CIN in fully developed gastric cancers. In line with the OGM analysis in organoids (**Fig. 1**), tumors with low *ARID1A* expression had a higher frequency of copy number (CN) gains, while CN losses were not significantly different (**Extended Data Fig. 1c**). Next, we evaluated the frequencies of genetic alterations of gastric cancer driver genes in eleven oncogenic signaling pathways in cancers with *ARID1A* loss. As expected, *ARID1A* deficiency was associated with increased missense mutations in these key cancer driver genes, given its previously reported function in regulating MMR (**Extended Data Fig. 1d**). Strikingly, the frequencies of SVs, including amplifications, deletions, and frameshift deletions, were more prominent than missense mutations in *ARID1A*-low tumors, supporting the role of ARID1A in regulating SVs. Together, these results from TCGA STAD suggest that the CIN phenotype induced by ARID1A deficiency at the premalignant stage is likely conserved in fully established cancer cells.

To confirm the causative effect of ARID1A deficiency on the CIN phenotype at the cancerous stage, we examined sister chromatid exchange (SCE) frequency, a widely used indicator of spontaneous CIN, in *ARID1A*-depleted cancer cells. We chose HCT116 cells as our model system for this assay because HCT116 is characterized as a CIN^-^ cell line with relatively high chromosomal stability ^23^. As shown in **Fig. 2b**, loss of ARID1A caused a significantly higher frequency of SCE. Notably, HCT116 is an MMR-deficient cell line. These data also suggested that the CIN phenotype induced by ARID1A depletion is independent of MMR status and is not restricted to gastric tissue type. Collectively, these results indicate that ARID1A deficiency induces characteristic features of CIN including chromosomal structural alterations and chromosome segregation errors, at both premalignant and cancerous stages, which set a stage of genomic instability to drive cancer development and evolution.

### ARID1A binds to α-satellite DNA and represses transcription of centromeric α-satellite RNA (α-SatRNA) through restricting RNAP II elongation

To investigate the molecular mechanisms underlying the CIN phenotype induced by *ARID1A* deficiency, we analyzed genome binding patterns of ARID1A using a previously published chromatin immunoprecipitation sequencing (ChIP-seq) dataset (GSE254865) in cancer cells ^24^. Surprisingly, we found that genes overlapping or near to *ARID1A*-bound repetitive sequences, such as transposable elements (TEs), were significantly enriched in the "mitotic cell cycle phase transition" pathway as annotated by GO analysis (**Fig. 2c**). We then further examined the binding of *ARID1A* to subclasses of repetitive sequences including long interspersed elements (LINEs), long terminal repeat elements (LTR), short interspersed elements (SINEs), and satellite DNAs. Among these repetitive sequences, ARID1A exhibited strong binding peak signals at satellite DNA compared to the genomic background (**Fig. 2d**). Specifically, ARID1A bound to both sense and antisense strands of centromeric α-satellite DNAs (**Fig. 2e**). We first conducted a ChIP-qPCR assay and verified that endogenous ARID1A was indeed bound to α-satellite DNAs (**Fig. 2f**). α-satellite DNAs are now known to undergo dynamic transcription catalyzed by RNA polymerase II (RNAP II) and generate α-satellite RNA (α-SatRNA), which plays an evolutionarily conserved role in maintaining chromosomal stability.

Next, we tested whether ARID1A regulates α-SatRNA transcription in a variety of models at different stages of tumor development (**Fig. 2g**). First, the qPCR results showed that *ARID1A* depletion induced α-SatRNA expression in premalignant gastric organoids (**Fig. 2g**). Second, *ARID1A* depletion or knockdown increased α-SatRNA expression in human gastric cancer cells and colon cancer cells respectively (**Fig. 2g**). Third, *ARID1A* knockdown promoted MajSatRNA and MinSatRNA expression (the murine counterparts of α-SatRNA) in a mouse gastric tumor cell line KPC, which was derived from a K-ras mutant; p53-knockout gastric genetic mouse model (**Fig. 2g**). Finally, to validate the clinical relevance of these findings, we conducted single molecule FISH (smFISH) with a probe set to examine α-SatRNA expression in peritoneal metastatic tumor specimens from gastric cancer patients. As we expected, low *ARID1A* expression was associated with higher α-SatRNA expression in gastric tumor cells (**Fig. 2h**).

To confirm that ARID1A is required for repressing α-SatRNA transcription and to further map the molecular regions of ARID1A necessary for this repression, we examined α-SatRNA expression in *ARID1A*-depleted cells reconstituted with WT or a series of *ARID1A*-deletion mutants. As shown in **Fig. 2i**, while WT *ARID1A* could restore the repression of α-SatRNA transcription, deletions in *ARID1A* (Δ 600-700 aa, Δ 1,500-1,600 aa, or Δ 1,600-1,700 aa) could not rescue increased α-SatRNA expression in *ARID1A*-depleted cells. These results led us to test whether these deletion mutants may impair the binding of ARID1A to α-satellite DNA. We found that ARID1A binding to α-satellite DNA is dependent on the region of 1,500-1,600 aa. In addition, *ARID1A* deletion of 600-700 aa, or 1,600-1,700 aa significantly impaired ARID1A binding to α-satellite DNA, albeit to a lesser extent than the 1,500-1,600 aa deletion, suggesting that multiple binding sites of *ARID1A* are required for its proper loading to α-satellite DNAs **(Fig. 2j)**. Uncontrolled or excessive expression of α-SatRNA is known to disrupt mitotic fidelity and result in CIN^25^. To demonstrate that ARID1A deficiency induces CIN via de-repression of α-SatRNA, we reconstituted *ARID1A* wildtype or deletion mutant (Δ 1,500-1,600 aa) in *ARID1A*-depleted HCT116 cells and conducted SCE assay. As expected, *ARID1A* wildtype, but not the deletion mutant (Δ 1,500-1,600 aa) can restore the frequency of SCE in *ARID1A*-KO cells **(Fig. 2k)**.

To further understand how ARID1A may regulate α-SatRNA transcription, we used ChIP-qPCR to examine total RNAP II occupancy on α-satellite DNAs. To our surprise, *ARID1A* depletion did not result in an increase in total RNAP II loading on α-satellite DNA regions **(Fig. 2l)**. In contrast, phosphorylated RNAP II at Serine 2 (pS2) was remarkably enriched on α-satellite DNAs in *ARID1A*-depleted cells compared to control cells **(Fig. 2l)**. Based on these findings, we reasoned that *ARID1A* depletion may increase the loading of CDK9 to induce excessive phosphorylation of RNAP II (pS2), which in turn facilitates RNAP II elongation and promotes α-SatRNA transcription. Indeed, we observed that *ARID1A* KO enhanced the binding of CDK9 to α-satellite DNAs **(Fig. 2l)**, suggesting a role of ARID1A in restricting CDK9-mediated RNAP II phosphorylation (pS2) and α-SatRNA transcription potentially via regulating RNAP II elongation. Together, these results show that ARID1A binds to repetitive DNA sequences of α-satellite DNA and is required for repressing α-SatRNA transcription. Loss of ARID1A induces the CIN phenotype potentially through unleashing α-SatRNA transcription via enhanced RNAP II elongation.

### PKMYT1i promotes α-SatRNA transcription and activates innate immune signaling in a α-SatRNA-dependent manner

The expression of α-SatRNA exhibits dynamic changes spatiotemporally during the cell cycle. It remains at a low level during G1/S but peaks at the G2/M transition, where it accumulates to prepare centromeres for mitosis^26^. Unlike many mRNAs and non-coding RNAs, α-SatRNA is not exported to the cytoplasm. However, during mitosis, α-SatRNA appears broadly distributed within the cytoplasm and is thus excluded from the nucleus when the nuclear envelope reforms^26^. Based on these observations, we hypothesized that the release of the G2/M cell cycle checkpoint may promote α-SatRNA transcription with enhanced sensitivity in ARID1A-depleted cells. The protein kinase, membrane-associated tyrosine/threonine 1 (PKMYT1) inhibitor (PKMTT1i) and the WEE1 kinase inhibitor (WEE1i) are promising G2/M checkpoint blockades under extensive clinical investigation, which remove the negative regulation of CDK1 kinase and thus drive the mitotic process ^27,28^. To test this hypothesis, we generated stable *ARID1A* knockdown or KO cells (**Fig. 3a**). As expected, in the gastric organoid model, PKMYT1i (RP6306) treatment selectively induced expression of α-SatRNA in *ARID1A*-KO cells (**Fig. 3b**). Similar results were observed in ARID1A-depleted human colon and gastric cancer cells treated with WEE1i (AZD1775) (**Fig. 3b**). MajSatRNA and MinSatRNA expressions in *Arid1a*-knockdown KPC mouse tumor cells were also significantly upregulated by AZD1775 treatment (**Fig. 3b**). In addition to quantitative analysis of α-SatRNA expression by RT-PCR, we used smFISH probe set to examine α-SatRNA expression and localization. PKMYT1i (RP6306) remarkably enhanced α-SatRNA expression in ARID1A-deficient cells and induced strong cytoplasmic signals of α-SatRNA (**Fig. 3c and 3d**).

**Fig 3.**
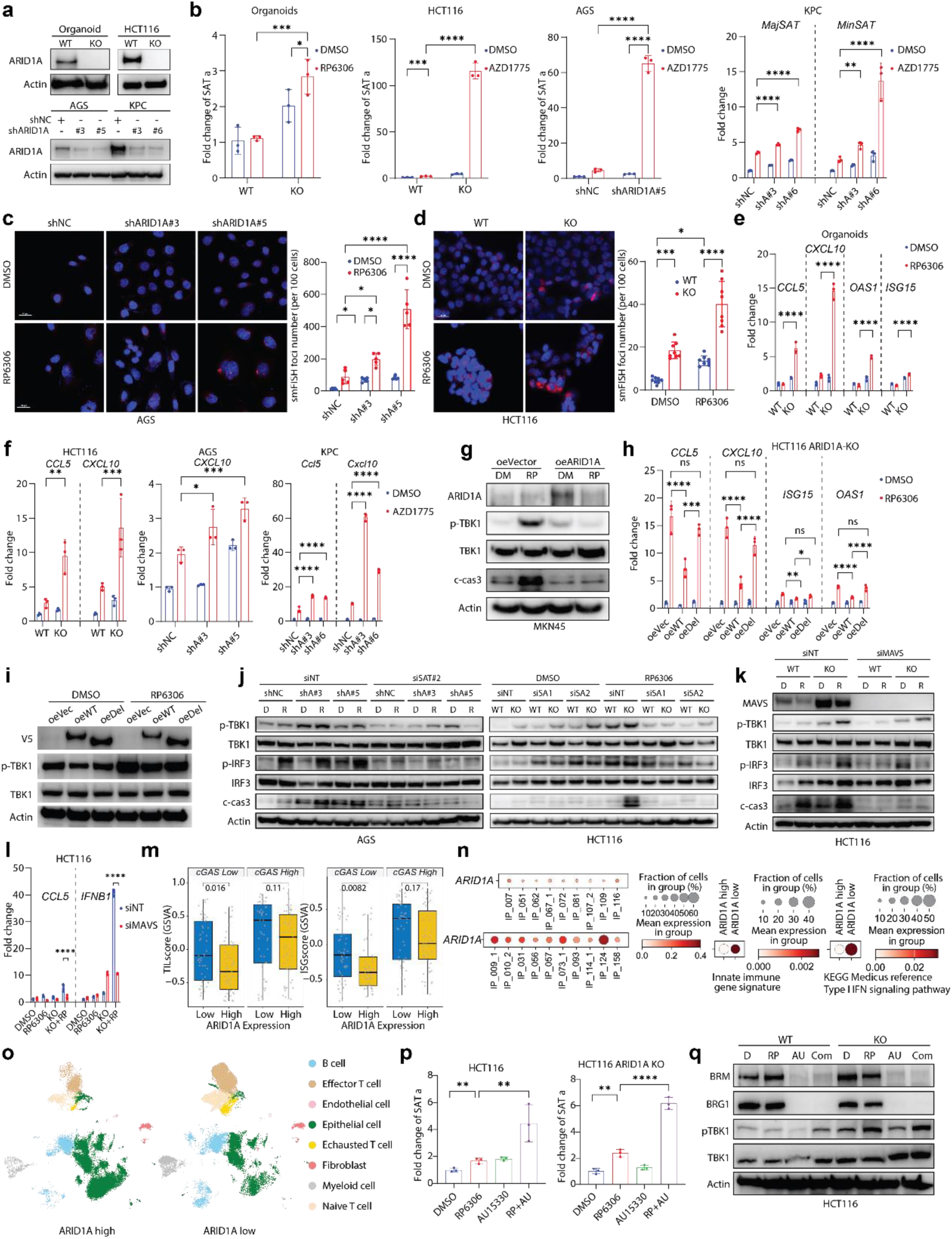
G2/M checkpoint inhibitors promote α-SatRNA expression and activate dsRNA-mediated innate immune signaling in *ARID1A*-deficient cells. **a,** Representative images of western blot (WB) for the knockout or knockdown efficiency of ARID1A in organoids, HCT116, AGS, ands KPC cells. **b,** qPCR results of α-SatRNA, MajSAT, and MinSAT in gastric organoids (RP6306 (2μM), 72h (n=3)), HCT116 (AZD1775 (0.5μM), 48h (n=3)), AGS (AZD1775 (1μM), 48h (n=3)), KPC (AZD1775 (1μM), 48h (n=3)) treated with indicated drugs. Significance was determined using two-way ANOVA. **c-d,** Representative images and quantification of smFISH staining for α-SatRNA in AGS (c, n=5) and HCT116 (d, n=8) cells treated with DMSO or RP6306 (2μM) for 48h. Two-way ANOVA was used. **e,** qPCR results showing the *CCL5*, *CXCL10*, *OAS1*, and *ISG15* changes in *ARID1A* WT or KO organoids treated with DMSO or RP6306 (2μM) for 72h (n=3). Two-way ANOVA test was used. **f,** qPCR results showing *CCL5* and *CXCL10* changes in HCT116 (left), AGS (middle), and KPC (right) cells treated with DMSO or AZD1775 for 48h (n=3). Two-way ANOVA was used. **g,** Empty vector (oeVector) or human *ARID1A* WT (oeARID1A) plasmids were transient transduced into MKN45 human gastric cancer cells, then cells were treated with DMSO or RP6306 (2μM) for 48h. WB was conducted to detect the ARID1A, phosph-TBK1 (p-TBK1), TBK1 and cleaved-caspase3 (c-cas3) protein expression. Representative images were shown. **h,** qPCR results of *CCL5*, *CXCL10*, *ISG15*, and *OAS1* in HCT116 ARID1A KO cells transduced with Vector (oeVec), *ARID1A*-WT (oeWT), or *ARID1A*-Δ 1500-1600 (oeDel) plasmids and treated with DMSO or RP6306 (2μM) for 48h. (n=3) Two-way ANOVA was used. **i,** WB of HCT116 *ARID1A* KO cells transduced with indicated plasmids and treated with DMSO or RP6306 (2μM) for 48h. Representative images were shown. **j,** *ARID1A* WT or knockdown/knockout AGS or HCT116 cells were transient transduced with siNT or siSAT (siSAT a#1 or #2), then cells were treated with DMSO or RP6306 (2μM) for 48h. WBs were conducted to detect the protein expression of phosph-TBK1 (p-TBK1), TBK1, phosph-IRF3 (p-IRF3), IRF3, and cleaved-caspase3 (c-cas3) in AGS (left) and HCT116 (right) cells. Representative images were shown. **k,** *ARID1A* WT or KO HCT116 cells were transient transduced with siNT or siMAVS, then cells were treated with DMSO or RP6306 (2μM) for 48h. WB was conducted to detect the protein expression of MAVS, phosph-TBK1 (p-TBK1), TBK1, phosph-IRF3 (p-IRF3), IRF3, and cleaved-caspase 3 (c-cas3). Representative images were shown. **l,** qPCR results of *CCL5* and *IFNB1* in HCT116 *ARID1A* WT or KO cells transduced with siNT or siMAVS and treated with DMSO or RP6306 (2μM) for 48h (n=3). Two-way ANOVA was used. **m,** GSVA analysis of the TILscore and ISGscore signature in ARID1A-low (25%, n=51) vs high (75%, n=51) groups within cGAS-low (50%, N=205) or high (50%, N=205) patients. STAD data was retrieved from TCGA database and student’s t test was used to determine the P value. **n,** Single-cell sequencing data from EGAS00001004443. Gastric cancer patients were divided into high (n=10) and low (n=9) two groups according to the ARID1A expression (left). Innate immune gene signature (middle) and KEGG Medicus reference type I IFN signaling pathway (right) were analyzed between the 2 groups. **o,** UMAP plot of indicated cell clusters in ARID1A-high and low groups. **p,** qPCR results of the α-SatRNA change in HCT116 *ARID1A* WT (left) or KO (right) cells treated with DMSO, RP6306 (2μM), or/and AU15330 (2μM) for 48h (n=3). One way ANOVA was used. **q,** WB results of the HCT116 cells treated with DMSO, RP6306 (2μM), or/and AU15330 (2μM) for 48h. Representative images were shown. *P<0.05, **P<0.01, ***P<0.001, ****P<0.0001.

Due to its repetitive, inverted sequence nature, and bidirectional transcription, α-SatRNA represents a major endogenous form of double-stranded RNA (dsRNA) derived from self-genome transcription. Next, we sought to determine whether the G2/M checkpoint blockades can enhance innate immune signaling via triggering the RNA sensing pathway in *ARID1A*-deficient cells. In gastric organoid models, PKMYT1i exhibited remarkable selectivity in inducing type-I interferon stimulated genes (ISG) such as *CCL5*, *CXCL10*, *OAS1,* and *ISG15* in *ARID1A*-depleted cells compared to control cells (**Fig. 3e**). Similarly, WEE1i led to a robust increase in the expression of *CCL5* and *CXCL10* in *ARID1A*-deficient cells in human cancer cell lines and *Arid1a*-KPC mouse gastric tumor cells (**Fig. 3f**). Moreover, we reconstituted *ARID1A* expression in the MKN45 gastric cancer cell line lacking endogenous ARID1A expression. As shown in **Fig. 3g**, the re-expression of ARID1A was sufficient to suppress the induction of p-TBK1 by PKMYT1i. It is worth noting that despite *ARID1A* deficiency enhancing innate immune signaling induced by the G2/M checkpoint inhibition in both premalignant organoid cells and cancerous cells, exclusive selectivity was observed in the *ARID1A*-depleted organoid model with very minimal effects of PKMYT1i observed in *ARID1A*-WT organoid cells. It is likely due to the relatively low level of genomic instability in these premalignant organoid cells without *ARID1A* depletion. In contrast, cancerous cells have higher levels of established genomic instability, which may render them sensitive to innate immune response and also cell death induced by PKMYTi/WEE1i due to loss of the G2/M checkpoint. Together, our data from premalignant and cancerous cell models of human and mouse origins demonstrated that ARID1A plays a determinant role in regulating innate immune signaling triggered by PKMYT1i/Wee1i regardless of different genetic contexts.

To further validate that enhanced activation of innate immune signaling in *ARID1A*-depleted cells in response to G2/M checkpoint blockade is dependent on α-SatRNA regulation, we conducted two sets of experiments. First, we reconstituted *ARID1A*-depleted cells with *ARID1A* wildtype construct or mutant construct lacking α-satellite DNA binding capacity. As shown in (**Fig. 3h and 3i**), while wildtype *ARID1A* re-expression significantly reduced PKMYT1i-induced *CCL5* and *CXCL10* expression, the mutant *ARID1A* without α-satellite DNA binding and SatRNA capacity restored activation of innate immune signaling. Second, we designed two specific siRNAs targeting the consensus α-SatRNA sequence (siSAT#1 and siSAT#2) as previously described^29^ to test whether the presence of α-SatRNA is required for PKMYT1i-induced innate immune signaling in *ARID1A*-KO cells. The knockdown efficiency of siRNAs against α-SatRNA was verified by qRT-PCR and smFISH staining in *ARID1A*-KO cells (**Extended Data Fig. 2a**). Silencing of α-SatRNA expression significantly blocked PKMYT1i-induced activation of TBK1 in both AGS and HCT116 cancer cell lines (**Fig. 3j**). Interestingly, PKMYT1i-induced activation of the apoptotic marker caspase-3 was also impaired by α-SatRNA knockdown (**Fig. 3j**). The knockdown efficiency of α-SatRNA at basal level and at the induced level by PKMIT1i was confirmed by qRT-PCR (**Extended Data Fig. 2b-2c**). Notably, α-SatRNA knockdown by siRNAs appeared to increase the basal level of p-TBK1 in both control and *ARID1A*-depleted HCT116 cells, but not in AGS cells (**Fig. 3j**), suggesting the basal level of p-TBK1 and PKMIT1i-induced p-TBK1 are likely regulated by different signaling inputs in HCT116 cells. Nevertheless, these data show that enhanced activation of key mediators of innate immune and apoptotic signaling (TBK1 and Caspas-3) induced by PKMYT1i is dependent on the presence of α-SatRNA.

### PKMYT1i amplifies MAVS-mediated dsRNA-sensing in ARID1A-deficient cancer cells

Previous study characterizing a series of colon cancer cell lines identified that HCT116 cells lack cGAS expression and are deficient in the cGAS-STING-mediated DNA sensing pathway^30^. Having determined that PKMYT1i-induced activation of innate immune signaling is dependent on α-SatRNA, we then examined whether PKMYT1i enhances innate immune response in ARID1A-deficient cancer cells via dsRNA-sensing mechanisms, particularly in HCT116 cells with pre-existing DNA sensing cGAS-STING pathway defects. The newly formed dsRNA is recognized and bound by OAS1 and subjected to RNase L for degradation. The degraded dsRNA can then be recognized by RNA sensors RIG1/MDA5, which in turn leads to the cascade activation of MAVS, TBK1/IRF3, and downstream ISGs ^31^. Thus, we first knocked down MAVS, a central molecule in the cytoplasmic dsRNA-sensing pathway. Reduced MAVS expression remarkably suppressed PKMYT1i-induced p-TBK1 and p-IRF3 expression, and consequently expression of ISGs including *CCL5* and *IFN-β* (**Fig. 3k and 3l**). Similar results were observed in AGS cancer cells (**Extended Data Fig. 2d and 2e**) and KPC mouse cells (**Extended Data Fig. 2f**). Additionally, we knocked down OAS1, encoding the 2’5’-oligoadenylate synthetase 1 (2-5A synthetase), that generates 2’, 5’-oligoadenylates and subsequently, activates RNase L and RNA sensors. As expected, *OAS1* knockdown significantly reduced activation of innate immune response in both HCT116 cells with cGAS-STING deficiency and AGS cells with competent cGAS-STING function (**Extended Data Fig. 2g and 2h**). These data suggest that MAVS-mediated dsRNA-sensing plays a critical role in mediating PKMYT1i-induced innate immune signaling in ARID1A-deficient cancer cells, regardless of the functional status of cGAS-STING pathway. This finding was further confirmed by the GSVA analysis that TILscore and ISGscore signatures in ARID1A low group were higher than those in the high groups within cGAS low gastric cancer patients (**Fig. 3m**).

To elucidate the clinical significance of these findings, we analyzed the scRNA-seq data of ascites cells from gastric cancer patients with peritoneal metastases and found that ARID1A deficiency correlated with an enriched innate immune gene signature and type I-IFN gene signature (**Fig. 3n**). Consistently, effector T cell infiltration was higher in the ARID1A-low group (**Fig. 3o**). In short, these data support a conventional cGAS-STING-DNA-sensing pathway-independent association of innate immune activation in ARID1A-deficient tumors.

Since *ARID1A* is an accessory subunit of the SWI/SNF chromatin-remodeling complex, where it cooperates with the catalytic ATPase subunits SMARCA2 or SMARCA4 to regulate chromatin accessibility and transcription, we further investigated whether SMARCA2/4 is involved in regulating α-SatRNA expression. We treated HCT116 cells with AU-15530, a proteolysis-targeting chimera (PROTAC) degrader of SMARCA2/4. In WT cells, AU-15530 treatment increased α-SatRNA expression at the basal level and further enhanced α-SatRNA expression induced by PKMYT1i in a similar manner to *ARID1A* depletion, while AU-15530 treatment did not further increase α-SatRNA expression in ARID1A-depleted cells (**Fig. 3p**). Furthermore, ARID1A depletion led to a 2-fold increase in α-SatRNA expression in the presence of PKMYT1i compared to WT cells (**Fig. 3p**). A similar fold increase in α-SatRNA expression (6-fold) was observed in ARID1A-depleted cells treated with PKMYT1i and AU-15530 compared to WT cells. Consistent with these findings, AU-15530 treatment enhanced PKMYT1-induced TBK-1 activation in WT cells while this increased TBK-1 activation was not observed in ARID1A-depleted cells (**Fig. 3q**). Taken together, these data suggest that the catalytic activity of the SWI/SNF complex is involved in regulating α-SatRNA transcription likely through ARID1A-mediated function.

### PKMYT1i selectively elicits cancer cell killing and immunomodulatory effects in ARID1A-deficient tumors

Based on these findings, we reasoned that PKMYT1i promotes α-SatRNA expression, exacerbates CIN and amplifies RNA sensing in *ARID1A*-deficient cancer cells, which may selectively kill and inflame these tumor cells to enhance therapeutic efficacy. To test this hypothesis, we first investigated the therapeutic effect of PKMYT1i using *ARID1A*-deficient human and mouse cell models. As expected, *ARID1A* depletion sensitized premalignant gastric organoids to PKMYT1i treatment by disrupting their structure and reducing organoid survival (**Fig. 4a**). This synthetic lethality was further confirmed across multiple assays in cancer cell lines, where PKMYT1i showed a dose-dependent reduction in cancer cell proliferation (**Fig. 4b**) and a significant increase in cell death assessed by apoptosis in *ARID1A*-deficient cells under a short treatment duration (**Fig. 4c**). During prolonged treatment, PKMYT1i decreased survival of *ARID1A*-deficient cancer cells as examined by clonogenic assay (**Fig. 4d**). Furthermore, we used mouse gastric tumor KPC cell line to test whether *Arid1a* deficiency enhances the therapeutic efficacy of PKMYT1i treatment as a monotherapy or in combination with immunotherapy in vivo using syngeneic mouse models. KPC tumor-bearing mice were randomly divided into 8 groups and were treated with vehicle, RP6306, anti-PD-L1 antibody, isotype IgG control, or combination treatment (**Fig. 4e**). Tumor volumes were measured every 2 days, and the endpoint of survival analysis was set as tumor size reaching 1000 mm^3^. In the group of *Arid1a* knockdown tumors treated with RP6306 plus anti-PD-L1 antibody, tumor volume showed the slowest growth rate (**Fig. 4f**). In addition, we conducted survival analysis to examine the overall therapeutic benefit. Despite the control groups receiving vehicle/IgG, *Arid1a* loss showed a positive effect on prolonging mouse survival, PKMYT1i and anti-PD-L1 antibody monotherapy significantly improved the survival of mice bearing Arid1a knockdown tumors (**Fig. 4g**). Most remarkably, the combination treatment of PKMYT1i and anti-PD-L1 antibody exhibited an exceptional therapeutic response in mice bearing *Arid1a* knockdown tumors. In this group, three out of five mice remained alive with very low tumor burden at the end of the experiment (100 days after tumor cell injection) compared to median survival time of 20 days for mice bearing Arid1a knockdown tumors in the absence of treatment (**Fig. 4g**). Furthermore, multiplex immunohistochemical (mIHC) staining of tumor samples from mouse models showed that *Arid1a* knockdown promoted CD8^+^ T cell infiltration and PD-L1 expression in either monotherapy or combination therapy groups (**Fig. 4h and 4i**). Collectively, these data show that ARID1A deficiency augments the anti-tumor efficacy of PKMYT1i therapy, particularly in combination with immunotherapy.

**Fig 4.**
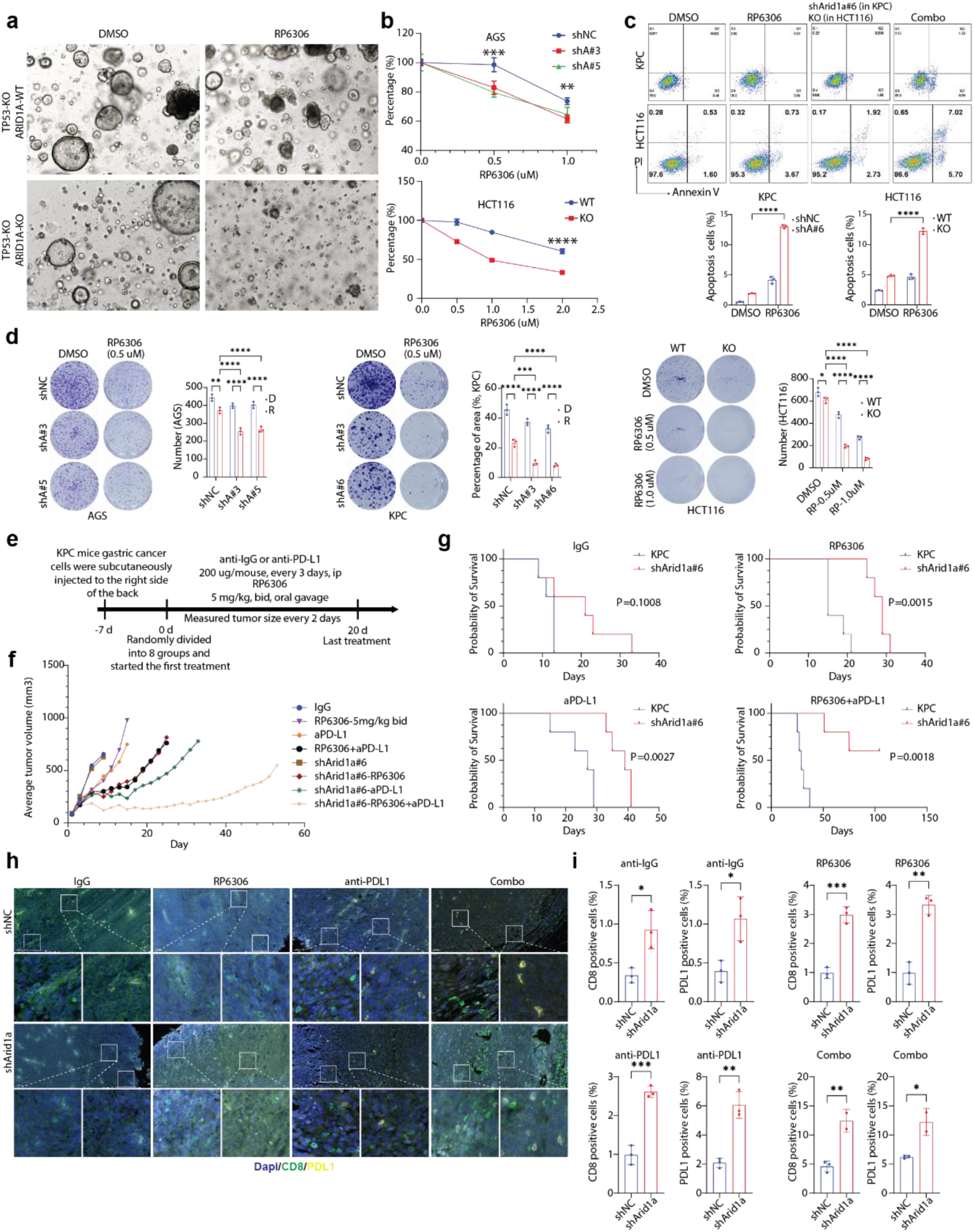
ARID1A deficiency enhances the anti-tumor effect of RP6306 in vitro and in vivo. **a,** Representative images of *ARID1A*-WT or -KO gastric organoids treated with DMSO or RP6306 (2μM) for 4 days. **b,** CCK8 assay of the AGS (upper) and HCT116 (lower) cells treated with DMSO or RP6306 in different concentration for 3 days. OD numbers were read at 450nm wavelength (n=5). Student’s t test or one-way ANOVA was used. **c,** Representative images and quantification of PI/AnnexinV staining flowcytometry in KPC and HCT116 indicated cells treated with DMSO or RP6306 (2μM) for 48h. Two-way ANOVA was used. **d,** Representative images and quantification of clony formation assay in AGS (left), KPC (middle), and HCT116 (right) cells treated with DMSO or RP6306 at the indicated concentration for 7 days (n=3). Two-way ANOVA was used. **e,** Scheme illustrating the mouse model construction and drug treatment timeline. **f,** Average tumor growth curve. Groups were shown as indicated. **g,** Survival curves of mice treated with anti-IgG, RP6306, anti-PD-L1, or combination (RP6306+anti-PD-L1) (n=5 in each group). Survival curve was drawn with the Kaplan-Meier method and compared using the Log-Rank test. **h,** Representative images of mIHC staining of CD8, PDL1, and DAPI in mice tissues. Scale bars were shown as 50 μm. **i,** Quantification of CD8- or PDL1-positive cells in anti-IgG, RP6306, anti-PD-L1, and combination (RP6306+anti-PD-L1) treatment groups. N=2 in the Arid1a knockdown combination treatment group while n=3 in the other groups. Significance was determined using student’s t test. *P<0.05, **P<0.01, ***P<0.001, ****P<0.0001.

## Discussion

Recent and continuously expanding genomic and single-cell analyses have unveiled an unexpectedly prevalent landscape of cancer-associated mutations and molecular signatures of CIN in normal cells and tissues^1–6,32–35^. What has been less clear is the pathobiological role played by cancer-associated mutations in normal tissues in promoting early CIN phenotypes and setting the stage for cancer to evolve. Here, we demonstrate that *ARID1A*, the most frequently mutated gene identified in both normal gastric epithelium and gastric cancer, is required for maintaining chromosome segregation and structural stability. Mechanistically, our study reveals a previously unknown role of the ARID1A-SWI/SNF chromatin remodeling complex in preventing the development of CIN at the precancerous stage by restricting RNAP II elongation at centromere repetitive satDNA sequences and thus limiting excessive α-SatRNA transcription. Furthermore, our findings provide a molecular framework linking aberrant α-SatRNA transcription in *ARID1A*-deficient cells to the activation of innate immune response mediated by the RIG1/MDA5/MAVS-dsRNA sensing pathway. Additionally, the dual molecular consequences of uncontrolled α-SatRNA on inducing CIN and dsRNA immune signaling can be further enhanced by disrupting the G2/M checkpoint, highlighting the potential of PKMYT1 inhibitor as an effective targeted therapy for patients with ARID1A-SWI/SNF-pathway deficient tumors or more broadly tumors with satRNAs overproduction.

Precise regulation of α-SatRNA transcription is not well known to play a critical role in maintaining chromosome fidelity during cell division^26^. Forcing overproduction of α-SatRNA in human and mouse cells causes chromosome segregation abnormalities, spontaneous DNA breaks and defects in DNA repair, which are key molecular changes leading to CIN^25,29,36–38^. For the first time, using human primary gastric organoids, our study has demonstrated that disrupted transcriptional control of noncoding centromere α-SatRNA in ARID1A-deficient cells may pave the way for the development of CIN at an early stage of tumorigenesis. This mechanistic connection between two fundamental functions of the ARID1A-SWI/SNF complex in gene transcriptional regulation and genome maintenance expands our understanding of how a cancer-associated mutation may create a molecular background of tolerable CIN in “normal cells” by disrupting proper transcription control of centromere repetitive DNA sequences. It was reported that the BRG1-BAF complex and the PBRM1-PBAF complex, two of the three mammalian SWI/SNF remodeling complexes, protect centromere integrity through regulating topoisomerase II alpha (TOP2A)-mediated DNA decatenation and establishing the heterochromatin marker H3K9 trimethylation (H3K9me3) at centromere and pericentromere regions ^39,40^. It is widely recognized that the SWI/SNF complex exhibits remarkable tissue specificity through the dynamic and combinatorial assembly of its diverse subunits ^41^. Aligned with our findings, distinct mechanisms governing the preservation of centromere stability may facilitate the adaptation of various subcomplexes to meet the specific requirements of cellular and tissue types, tumor development stages, and cell division stimuli. Previous studies from our group and others showed impaired MMR repair and MSI in *ARID1A*-deficient cancer cells ^19,20,42^. Notably, our data demonstrated that ARID1A-deficient gastric organoids exhibited molecular alterations of CIN without significantly increasing mutation burden or compromising essential MMR factors. These data indicate that the two primary types of genomic instability, CIN and MSI, may manifest at distinct stages of tumor development. We speculate that the additional molecular and cellular prerequisites necessary for the heightened mutation burden or MSI observed in ARID1A-deficient cancer cells may not be present in gastric organoids at the precancerous stage.

Innate immune signaling initiated by CIN or, more broadly, by DNA damage and genomic instability is primarily attributed to the formation of micronuclei and the subsequent release of double-stranded DNA (dsDNA) into the cytoplasm ^43,44^. This dsDNA activates the cGAS-STING-mediated innate immune response ^43,44^. Consistent with this notion, a recent study reported that *ARID1A* depletion increases R-loops, a DNA:RNA hybrid with a displaced strand of DNA, induces cytosolic single-stranded DNA (ssDNA), activates STING-dependent type I IFN signaling and thus improves immune responsiveness of cancer cells^45^. Our study demonstrates that aberrant expression of α-SatRNAs in ARID1A-deficient cells not only triggers CIN, but also elicits type I IFN signaling through the RIG/MDA5/MAVS-dependent dsRNA sensing pathway in a cGAS-STING DNA sensing pathway-independent manner. These findings suggest a previously uncharacterized role of dsRNAs derived from α-SatRNA transcripts in inducing inflammation-associated CIN and synergistically fueling cancer evolution. It is plausible that two distinct RNA and DNA mechanisms, mutually reinforcing, initiate type I IFN signaling and synergistically promote innate immune responses in *ARID1A*-mutant tumors.

Recent findings indicate that a generalized mRNA vaccine, such as the COVID-19 mRNA vaccine, without targeting specific tumor proteins could enhance cancer immunotherapy, suggesting the significance of engaging RNA sensing pathways in immunotherapy ^46^. Our findings reveal that aberrant transcription of α-SatRNAs can serve as a novel source of self-dsRNA derived from the noncoding genome, thereby modulating the innate immune response. In contrast to many self-derived RNAs including LINEs, SINEs and endogenous retrovirus (ERVs) ^47^, α-SatRNAs typically are not exported to the cytoplasm and predominantly localized in the nucleus^26^. However, during all stages of mitosis, α-SatRNAs are broadly distributed in the cytoplasm and thus excluded from the nucleus when the nuclear envelope reforms^26^. On the basis of this observation, we reasoned that disrupting G2/M checkpoints may facilitate expression and release of α-SatRNAs into the cytoplasm by inducing premature and uncontrolled mitosis. This process could lead to enhanced cell death and dsRNA sensing-mediated innate immune response in cancer cells.

Indeed, our data demonstrates that *ARID1A-SWI/SNF* deficiency and/or aberrant α-SatRNAs expression in cancer cells serves as a novel dual target for PKMYT1i. On one hand, PKMYT1i induces cell death by disrupting the G2/M cell cycle checkpoint and aggravating the molecular consequences of CIN in these cells. On the other hand, PKMYT1i further promotes aberrant expression of α-SatRNAs and the release and engagement of these self-dsRNAs with RNA sensors, such as RIG-I/MDA5/MAVS, which mediate innate immune signaling and thus enhances the efficacy of immunotherapy. The prevailing therapeutic strategy for G2/M cell cycle checkpoint blockades targeting PKMYT1, which function as negative regulators of CDK1, is predicated on the concept of synthetic lethality when combined with CCNE1 amplification in cancer cells. Based on this model, the defined key vulnerability for PKMYT1i/WEE1i is CCNE1 amplification associated with replication stress and mitotic catastrophic-induced cell death ^27,28,48^. Our study elucidates a novel mechanistic rationale employing PKMYT1/WEE1 inhibitors to target tumors exhibiting ARID1A-SWI/SNF deficiency or, more broadly, tumors harboring aberrant α-SatRNA expression as monotherapy or in conjunction with immunotherapy. Notably, the inactivation of cGAS/STING is prevalent in a variety of human cancers ^30,49–51^. This novel therapeutic paradigm holds significant importance in guiding the selective activation of innate immune signaling through the α-SatRNAs-dsRNA innate immune signaling axis by PKMYT1i/WEE1i in tumors that have pre-existing inactivation or silencing of DNA sensing pathways. Additionally, pharmacological approaches have been developed and tested in clinical studies to disrupt SWI/SNF function by catalytic inhibition and proteolysis-targeting chimera (PROTAC)-mediated degradation of SMARCA2 (BRM) and SMARCA4 (BRG1) subunits with ATPase activity ^52^. Our data demonstrate that chemically degrading BRG1/BRM induces the expression of α-SatRNAs. This mechanistically supports the requirement for ARID1A-SWI/SNF chromatin remodeling activity in regulating α-SatRNA transcription, and indicates a possibility of chemically inducing aberrant α-SatRNA-based therapeutic vulnerability in cancer cells without genetic alterations in the ARID1A-SWI/SNF complex. Furthermore, the quantitative detection of α-SatRNAs by RNA-FISH analysis can be employed as a clinically applicable biomarker for PKMYT1/WEE1 inhibitors or in conjunction with immunotherapy. This warrants further investigation utilizing clinical samples.

## Methods

### Cell culture

Human gastric cancer cell lines AGS (RRID: CVCL_0139) and MKN45 (RRID: CVCL_0434) were provided by Jaffer A. Ajani’s laboratory. Mouse gastric tumor cell line KP-Luc2 (KPC) was provided by Jaffer A. Ajani’s laboratory. Human colon cancer cell line HCT116 WT and *ARID1A* -KO (Q456+/Q456+) were purchased from Horizon Discovery Ltd. Human gastric cancer cell lines were cultured in RPMI-1640 medium (Corning) and the mice cell lines were cultured in DMEM medium (high glucose, Corning). HCT116 cells were cultured in McCoy’s 5A medium (10-050-CV, Corning). All mediums were supplemented with 10% FBS.

Human gastric organoids (ARID1A^WT^/TP53^KO^, ARID1A^KO^/TP53^KO^) were provided by Yuan-Hung Lo’s laboratory and cultured with 50% WENR medium^18^. All cells and organoids were incubated in a humidified 37°C incubator with 5% CO2.

### Reagents, kits and antibodies

The PKMYT1 inhibitor RP6306 (HY-145817A), SMARCA2/4 degrader AU-15330 (HY-145388), and ROCK inhibitor (HY-10071) were purchased from MedChemExpress (RRID: SCR_025062). Cell Counting Kit-8 (CK04-11) was purchased from Dojindo Molecular Technologies, Inc. The reagents for single molecular fluorescence in situ hybridization (smFISH) including the Stellaris® RNA FISH Hybridization Buffer (SMF-HB1-10), Wash Buffer A (SMF-WA1-60), Wash Buffer B (SMF-WB1-20), and probe sets were purchased from LGC Biosearch Technologies. The SimpleChIP® Enzymatic Chromatin IP Kit (#9003) was purchased from Cell Signaling Technology (CST, RRID: SCR_002071).

For antibodies used in this paper, anti-ARID1A (#12354, RRID:AB_2637010), phospho-TBK1/NAK (#5483, RRID:AB_10693472), TBK1 (#3504, RRID:AB_2255663), phospho-IRF3 (#29047, RRID:AB_2773013), IRF3 (#4302, RRID: AB_1904036), cleaved Caspase-3 (#9661, RRID: AB_2341188), phospho-Histone H2A.X (#9718, RRID:AB_2118009), MAVS (#24930, RRID: AB_2798889), Brg1 (#49360, RRID: AB_2728743), BRM (#11966, RRID: AB_2797783) were purchased from Cell Signaling Technology (CST). And the beta Actin Monoclonal Antibody (# AM4302, RRID: AB_2536382) was purchased from Invitrogen. The HRP conjugated second antibodies anti-rabbit IgG (#7074, RRID: AB_2099233) and anti-mouse (#7076, RRID: AB_330924) were purchased from CST.

### Plasmids, lentivirus and siRNA

The V5 tagged human ARID1A full length (FL), truncated plasmid that delete the 1500-1600 residues (del1500-1600) were generated previously using the corresponding Vector and kept in our lab. The H2B-mCherry overexpression plasmid was a gift from Robert Benezra (Addgene plasmid # 20972; http://n2t.net/addgene:20972; RRID: Addgene_20972).

Lentivirus packaging was conducted as described previously. The plasmids that targeting human ARID1A or mouse Arid1a were purchased from Sigma-Aldrich (RRID:SCR_008988). And the following sequences were used: shARID1A#3-CGGCTCACAATGAAAGACATT, shARID1A#5-GCCTGATCTATCTGGTTCAAT, shArid1a#3-CCCAGAGATTGGTCTTGGAAA, shArid1a#6-CCTAGGCAGCCTAACTATAAT.

siRNA was used for transient knockdown in this study. The pre-designed siRNA set targeting human MAVS (HY-RS08176) or mouse Mavs (HY-RS17037) was purchased from MedChemExpress. The Dharmacon on-target siRNA pool that targeting human OAS1 was purchased from Horizon Discovery. The siRNA targeting satellite RNA a was synthesized by Sigma-Aldrich and the sequence were as following: siSAT-a#1_ ACGGGAAUAUCUUCAUAUAAA, siSAT-a#2_ GGAAGGUUCAACUCUGUUACU. The siRNA Universal Negative Control (siNT, # SIC001) was purchased from Sigma-Aldrich.

### Sister chromatid exchange (SCE) assay

The HCT116 WT and ARID1A-KO cells were cultured in 10 cm dishes and were treated with RP6306 for 36h, followed by colcemid (0.1 ug/ml) for 2h. The cells were harvested, gently resuspended in 8 ml pre-warmed 0.075M KCL and incubated at 37°C for 10min. After centrifugation, cells were pre-fixed with cold Carnoy’s solution (3:1 methanol:glacial acetic acid), followed by further fixed twice. Finally, cells were resuspended in 2ml Carnoy’s solution.

Cell samples were dropped to the cold humidified slides and dried at room temperature. The slides were incubated with Hoechst 33258 solution (150ug/ml of ddH20) for 15 min in dark room at room temperature. After washed with distilled water, the slides were put into the 2x SSC buffer and exposed to UV light for 2 hours. Then the slides were incubated in 2x SSC buffer at 65°C for 1.5h. The slides slowly cooled down at room temperature and washed with distilled water, then stained with 5% Giemsa for 3 min. The chromatin was captured using a Zeiss microscope and analyzed by CytoVision system (Leica Microsystems Inc, RRID:SCR_025697).

### Clonogenic assay and CCK8 assay

For the clonogenic assay, 800 cells were plated to the 6-well plate for 24h prior to the treatment. After 5-10 days treatment with the indicated reagents, colonies were fixed with 4% PFA for 30 min at room temperature, then stained with 0.1% (w/v) crystal violet solution for 15 min. Cell colonies were photographed and counted after washing.

For the CCK8 assay, cells were seeded in the 96 well plate (1500 cells per well). Indicated treatment were added the next day and lasted for 3 days. Then add 10 uL CCK8 solution to each well of the plate, incubated in 37℃ for 30 min and then read the absorbance at 450 nm using a microplate reader (RRID: SCR_019748).

### PI/AnnexinV staining flow

HCT116 or KPC cells were treated with the indicated drugs for 48h, then cells were harvested with non-EDTA trypsin. After washing with PBS twice, cells were resuspended in AnnexinV binding buffer and stained with AnnexinV and PI for 15min. Then cells were subjected to flow cytometry analysis. The apoptosis cell numbers and the proportion were counted.

### Western blot

After transfected with indicated siRNA and treated with drugs, cells were harvested and lysed with urea buffer (8M urea, 50mM Tris-HCl pH8.0, 10mM β-mercaptoethanol, 1x protease inhibitor cocktail, 1x phosphorylated protease inhibitor). Protein concentration was measured with the bradford solution (#5000006, Bio-Rad). Total cellular lysates were subjected to 4–20% gradient SDS-PAGE and transferred to an Immobilon-P PVDF membrane (IPVH00010, Merck Millipore). The membrane was blocked with 5% BSA in 1x TBS-T (0.01 M Tris base, 0.15 M NaCl, 0.05% Tween-20). The aforementioned primary antibodies were used for western blotting, anti-Actin was diluted at 1:5000, and the other primary antibodies were diluted at 1:1000. After washing with TBS-T, the membranes were incubated with the corresponding secondary antibodies (1:3000). The western blots were visualized using Clarity Western ECL substrate (Bio-Rad 170–5061) and scanning with a ChemiDocMP imaging system (Bio-Rad).

### Quantitative real-time PCR (qPCR)

Total RNAs (1ug) were purified with PureLink RNA mini kit (#12183018A, Thermo Fisher Scientific), then reverse transcribed into cDNA by using HiScript IV RT SuperMix (#R423-01, Vazyme). SsoAdvanced Universal SYBR Green Supermix (#1725274, Bio-Rad) was used for the thermocycling reaction in an Applied Biosystems ViiA 7 Real-Time PCR System (RRID:SCR_023358). The primers used in this study were shown in supplementary table1.

### Single molecular FISH (smFISH) of RNA

The smFISH was conducted according to the manufacturer’s protocol. For cultured cells, grow cells on 18 mm coverslip in 12 well plate. After treated with indicated drugs, aspirated the medium and washed with PBS. Then added 1 ml fixation buffer (3.7% (vol./vol.) formaldehyde in 1X PBS) and incubated at room temperature for 10 min. After fixation, wash the coverslip with PBS twice. Then immersed the slides in 70% ethanol for 4h at 2∼8°C to permeabilize. Aspirated the 70% ethanol and incubated the cells with buffer A (90% wash buffer A + 10% formamide) for 5 min at room temperature. And cells were incubated with the α-SatRNA probes mix (1.25 uM probes, 10% formamide, and 90% hybridization buffer) in a humidified chamber at 37 °C overnight. The next day, removed the probes mix and incubated with fresh buffer A in the dark at 37 °C for 30 min. Then washed with buffer B at room temperature for 5 min. Mounted the coverslips with ProLong™ Gold Antifade Mountant with DNA Stain DAPI (P36931, Thermo Fisher Scientific) and subjected to whole slides scan by using Akoya Biosciences Vectra 3 Automated Quantitative Pathology Imaging System (RRID:SCR_025828).

For paraffin-embedded (FFPE) gastric cancer tissues, sections were sliced at a thickness of 5 um. After deparaffinization, the slides were incubated with pre-warmed (37 °C) proteinase K solution (10 μg/mL proteinase K in 1X PBS) for 20 min at 37 °C. Then the slides were subjected to hybridization with the same steps of the cultured cells. Finally, the slides were scanned by the Vectra 3 System (RRID:SCR_025828). The probe set sequences were shown in supplementary table2.

### Live cell image

Live cell imaging was conducted in the University of Texas MD Anderson Cancer Center Advanced Microscopy Core Facility (RRID:SCR_026611). For the live cell imaging, organoids were transfected with the lentivirus containing an H2B-mCherry construct (RRID: Addgene_20972). Before the imaging, these organoids were dissociated using TrypLE Express Enzyme (Gibco, #12604021) and plated in 12μl Matrigel (Corning, #356231) in a Lab-Tek II Chambered Coverglass (Thermo Fisher Scientific, #155409). Three days later, the chamber was subjected to a Leica SP8 laser-scanning confocal microscope (RRID:SCR_018169), equipped with atmospheric and temperature control. H2B-mCherry positive organoids were imaged in xyzt mode for 20-24h at 5 min intervals using a ×63 oil objective. In total, 9-33 z-sections were imaged per organoid. Raw data were processed by the Leica Application Suite X (RRID:SCR_013673) software and converted to videos using ImageJ (RRID:SCR_003070). Z-sections were separated and stained with different colors using the “Fire” panel. Several organoids were observed in each group, the normal and abnormal chromosome segregations in each organoid were counted and the abnormal proportion was calculated.

### Chromatin Immunoprecipitation (ChIP) qPCR

ChIP qPCR was used to study the proteins that bind to the α-Sat DNA and verify the binding domain. The assay was conducted with the SimpleChIP Kit (#9003S, Cell Signaling) and according to the manufacture’s protocol. Briefly, chromatin-protein was crosslinked and the cells were harvested. After digested with micrococcal nuclease, the lysate was subjected to sonication. The DNA was broken into 150–900 bp fragments. For each IP reaction, 10 ug DNA was used, the indicated antibodies were incubated with the samples at 4 °C overnight followed by further incubated with magnetic beads for 4h. The DNA products were quantified by qPCR, and the reaction system was 10 µL (2 µL DNA product, ∼50 ng). Quantitative results were displayed as enrichment to input (percent input = 2% × 2^(CT^ ^2%^ ^input^ ^sample – CT IP sample)^) or corresponding fold change. Anti-rabbit IgG or anti-mouse IgG were used as a negative control. The qPCR primers used are: α-SatRNA-F: AAGGTCAATGGCAGAAAAGAA; α-SatRNA-R: CAACGAAGGCCACAAGATGTC.

### Bioinformatic analysis

The RNA seq data of the WT and ARID1A KO organoids were retrieved from GSE164179. The differentially expressed genes (DEGs) were defined as FDR<0.05 and Log2Foldchange>1 according to the analysis results by ‘DESeq2’ (RRID:SCR_015687) package. The ‘clusterProfiler’ package (RRID:SCR_016884) was applied to analyze the enriched pathways.

The Cancer Genome Atlas (TCGA, RRID:SCR_003193) database was used to analyze the DEGs and pathways related to ARID1A. TCGA-STAD data was downloaded using the ‘TCGAbiolinks’ package (RRID:SCR_017683). According to the ARID1A expression, the top 25% patients were defined as ARID1A high groups and the low 25% was defined as ARID1A low group. The ‘DESeq2’ package (RRID:SCR_015687) was used to analyze the DEGs while the ‘clusterProfiler’ package (RRID:SCR_016884) was applied to analyze the enriched pathways. For the gene signature analysis, the ‘GSVA’ package (RRID:SCR_021058) was used. The TIL score was defined as a gene set of 18 genes and the ISG score was defined as a 38-genes set. The gene lists were shown in Supplementary table3. These analyses were conducted in R software (version 4.4.1, RRID:SCR_001905).

To analyze the repeat sequence that ARID1A binding to, the ARID1A ChIP data (GSE254865) was downloaded from Gene Expression Omnibus (GEO) (RRID:SCR_005012). The human RepeatMasker (RRID:SCR_012954) annotation (version AH99003) was downloaded by the ‘AnnotationHub’ package (RRID:SCR_024227). The peaks were re-annotated and the most frequent repClass type was defined as the main type for the peak. The proportion of different repeat sequence were calculated individually in the ChIP data and genome background, then the enrichment score was calculated. The enrichment heatmap for the centromere satellite DNA was made with the ‘EnrichedHeatmap’ package (RRID:SCR_023082). To analyze the repeat sequence related genes pathway, these peaks were annotated using the ‘TxDb.Hsapiens.UCSC.hg38.knownGene’ package. The transposable elements corresponding genes were identified and then subjected for the enrichment analysis using the ‘clusterProfiler’ package (RRID:SCR_016884).

The single cell RNA sequence data analysis was conducted as previously described. The data were retrieved from EGAS00001004443. The Cell Ranger Software (RRID:SCR_017344) was used for the raw data process, and the “Seurat” package (RRID:SCR_016341) was used for the normalized UMI count data. The 19 gastric cancer patients were divided into 2 groups according to ARID1A expression, and several gene signatures and the infiltrated immune cells in the two groups were compared.

### Multiplex immunohistochemistry (mIHC) staining

The mIHC staining for the tumor tissues were conducted using the Opal™ 4-Color anti-Rabbit Manual IHC Kit (NEL840) according to the manufacture’s instructions. Briefly, paraffin embedded tissues were sectioned with the thickness of 5um. After deparaffined and rehydrated, slides were fixed with 10% neutral buffered formalin. Then slides were subjected to antigen retrieval using the microwave methods followed by blocking. The primary antibody was incubated overnight at 4°C, and the anti-rabbit secondary antibody was incubated at room temperature for 10 min. After the opal signal was generated, the slides were subjected to microwave treatment again and followed by the new round of staining. The primary antibodies used in the mIHC were anti-CD8 (CST, #98941, 1:200, RRID:AB_2756376), anti-PD-L1 antibody (CST, #64988, 1:200, RRID:AB_2799672). Finally, the slides were mounted with ProLong™ Gold Antifade Mountant with DNA Stain DAPI (ThermoFisher, P36931) and subjected to scan by the Akoya Biosciences Vectra 3 Automated Quantitative Pathology Imaging System (RRID:SCR_025828).

### Optical genomic mapping (OGM)

The OGM assay was conducted by the MDACC Advanced Technology Genomics Core Facility (RRID:SCR_026261). Briefly, organoids were dissociated and filtered using a 70 μm Sterile Cell Strainers (Fisher Scientific, # 22-363-548). Totally 5 million single cells per group were prepared. The cells were then used for isolating ultra-high molecular weight DNA with a minimal size of 150 kb, using the Bionano Prep SP-G2 Blood and Cell DNA Isolation Kit (#80060). The DNA was homogenized and measured the concentration. Then with a single enzymatic reaction using the Direct Label and Stain (DLS) Kit (#80005), DNA was labeled and homogenized. The labeled DNA was loaded into the chip (#20440) and subjected to the Saphyr system (RRID: SCR_017992) for the scanning. About 100X of raw data was collected and structure variant analysis was performed to generate the curated list of annotated variants.

### WES sequence

The WES sequence of organoids was conducted by Novogene Co, Ltd.

### In vivo mouse models

For the in vivo mouse models, 5×10^5^ KPC shNC or shArid1a gastric mouse tumor cells were injected subcutaneously to the right side of C57BL/6 mice (male, 5-7 weeks old). One week later, the mice were randomly assigned to different groups. Tumor bearing mice treated with isotype IgG (i.p.) or anti-PD-L1 antibody (i.p. 200ug/mouse, Bio X Cell Cat# BE0361, RRID: AB_2927503) every 3 days, RP6306 (5mg/kg, oral delivery) twice per day. The total treatment time was 21 days. The tumor size and mice body weight were measured every 2 days, and tumor size was calculated as the (length x width^2^)/2. Mice reaching a humane endpoint that tumor size more than 1000 mm^3^ were euthanized. The survival time was recorded. There were 5 mice in each group. The mouse study was conducted in compliance with protocol approved by the MD Anderson Cancer Center Institutional Animal Care and Use Committee.

### Statistical analysis

GraphPad Prism 10.3 (RRID:SCR_002798) was used for statistical analysis in this study. Student’s t-test was used to compare mean value between two groups while one-way or two-way ANOVA were used to analyze the differences between multiple groups. Statistical significance was set at two-tailed *P* < 0.05 in this study.

---

**Extended Data Fig. 1.**
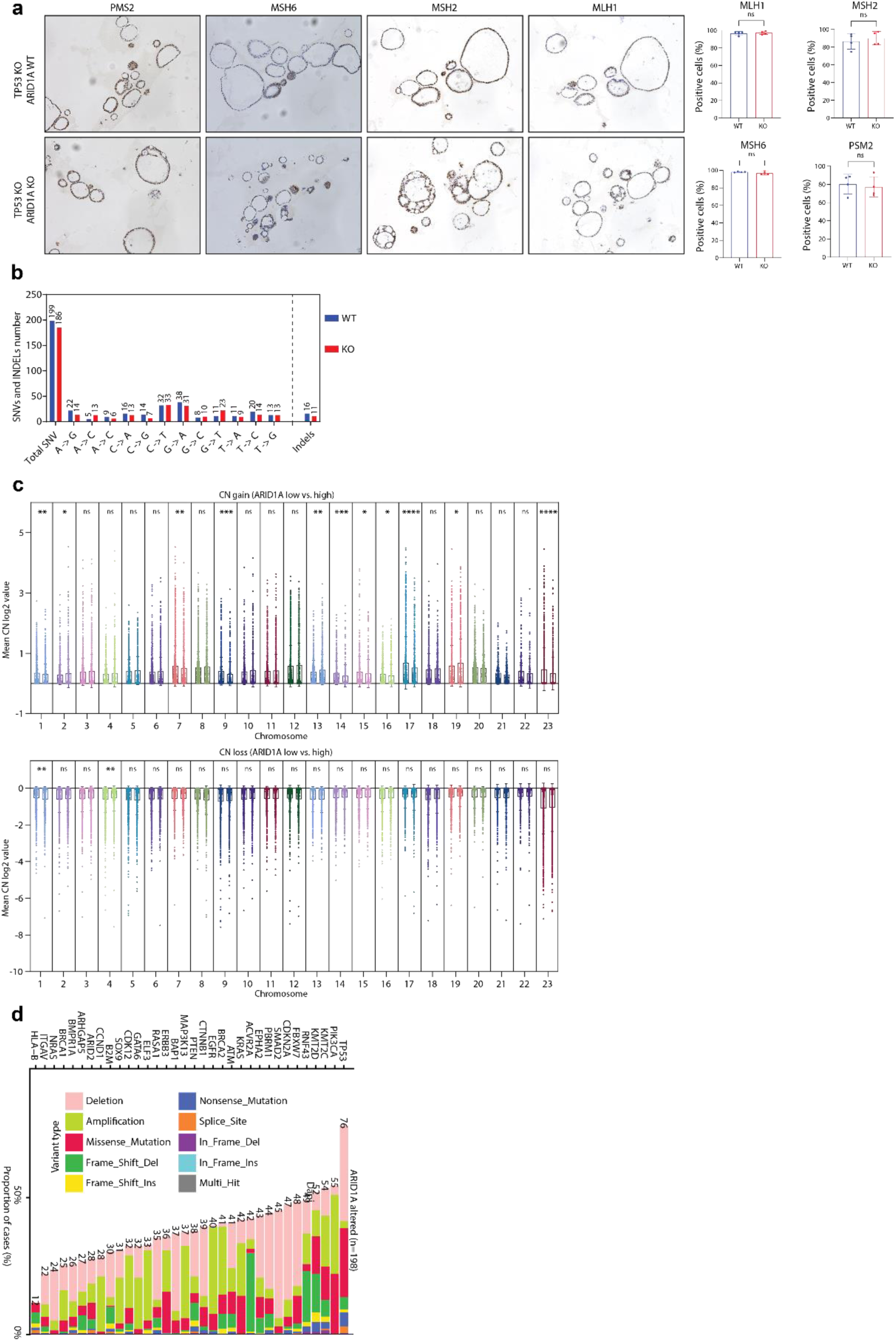
ARID1A deficiency impairs chromosomal stability but not MMR at the premalignant stage of gastric tumorigenesiss. **a,** Representative images and quantification of IHC staining for PMS, MSH6, MSH2, and MLH1 in ARID1A WT or KO gastric organoid (n=4). Student’s t test was used. **b,** Quantification of the single nucleotide variant (SNV) and indels in ARID1A WT or KO gastric organoids, determined by WES. **c,** Copy number gain (upper) and loss (lower) comparison between ARID1A-low and -high gastric cancer patients from the TCGA database. Significance was determined by student’s t test. **d,** CNV and mutations of the indicated genes in gastric cancer patients with ARID1A mutation or deletion (n=198). TCGA STAD cohort was used. ns. no significance. *P<0.05, **P<0.01, ***P<0.001, ****P<0.0001.

**Extended Data Fig. 2.**
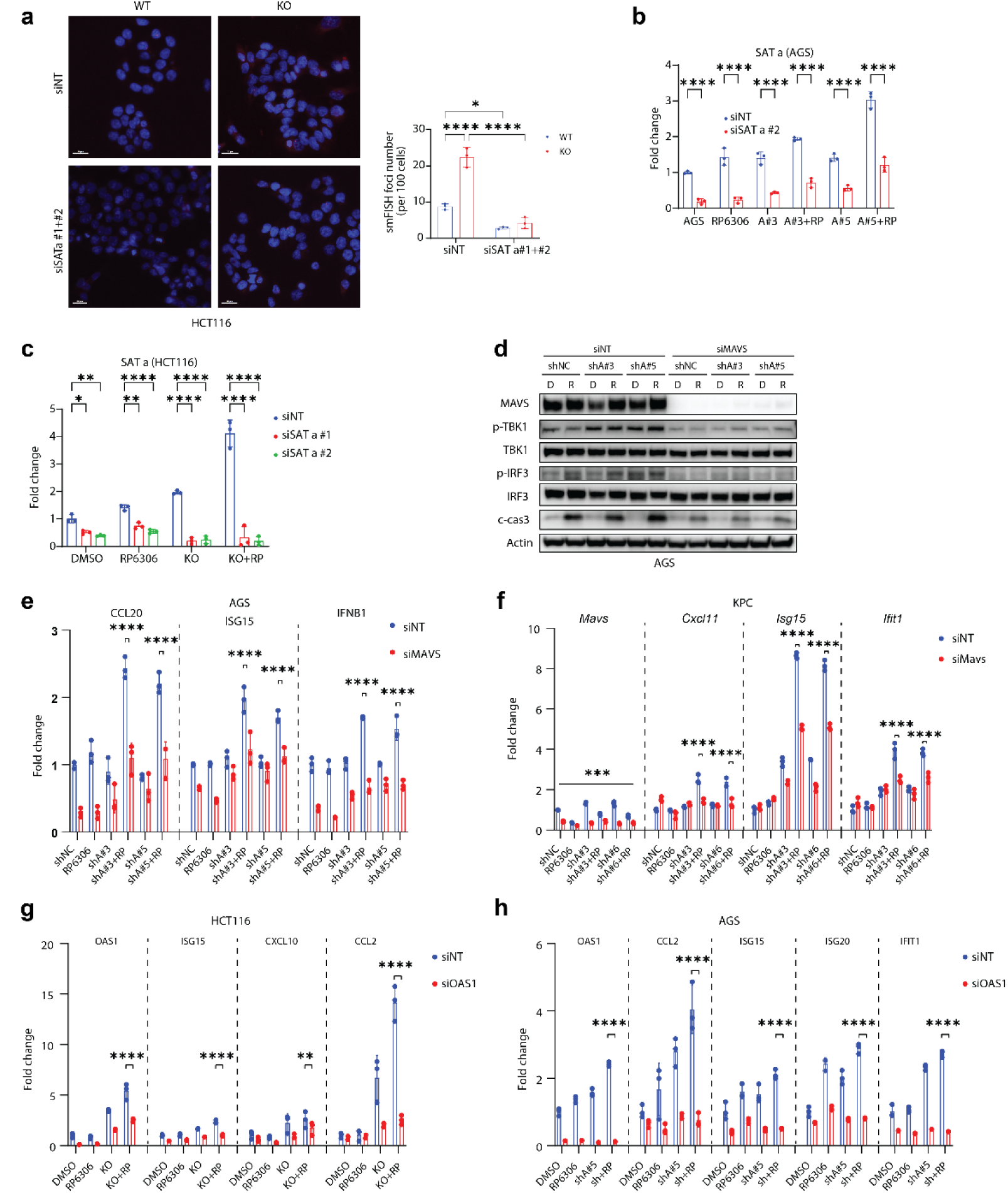
RP6306 promotes innate immune response in *ARID1A*-deficient cells via α-SatRNA. **a,** Representative images and quantification of smFISH staining in HCT116 *ARID1A*-WT or KO cells transduced with siNT or siSAT #1/#2 (n=3). Two-way ANOVA was used. **b-c,** *ARID1A* WT or knockdown/knockout AGS or HCT116 cells were transiently transduced with siNT or siRNA targeting α-SatRNA (siSAT#1 or #2), then cells were treated with DMSO or RP6306 (2μM) for 48h. qPCR was conducted to verify the knockdown efficiency of α-SatRNA in AGS (b) and HCT116 (c) cells (n=3). Two-way ANOVA was used. **d,** *ARID1A* WT (shNC) or knockdown (shARID1A #3 and #5) AGS cells were transiently transduced with siNT or siMAVS, then cells were treated with DMSO or RP6306 (2μM) for 48h. WB was conducted to detect the protein expression of MAVS, phosph-TBK1 (p-TBK1), TBK1, phosph-IRF3 (p-IRF3), IRF3, and cleaved-caspase3 (c-cas3). Representative images were shown. **e,** *ARID1A* WT (shNC) or knockdown (shARID1A #3 and #5) AGS cells were transiently transduced with siNT or siMAVS, then cells were treated with DMSO or RP6306 (2μM) for 48h. qPCR was used to detect the mRNA changes of CCL20, ISG15, and IFNB1 (n=3). Two-way ANOVA was used. **f,** Arid1a wt (shNC) or knockdown (shArid1a #3 and #6) KPC mouse gastric cancer cells were transiently transduced with siNT or siMavs, then cells were treated with DMSO or RP6306 (2μM) for 48h. qPCR was used to detect the mRNA changes of Mavs, Cxcl11, Isg15, and Ifit1 (n=3). Two-way ANOVA was used. **g,** *ARID1A* WT or KO HCT116 cells were transiently transduced with siNT or siOAS1, then cells were treated with DMSO or RP6306 (2μM) for 48h. qPCR was used to detect the mRNA changes of OAS1, ISG15, CXCL10, and CCL2 (n=3). Two-way ANOVA was used. **h,** *ARID1A* WT (shNC) or knockdown (shARID1A#5) AGS cells were transiently transduced with siNT or siOAS1, then cells were treated with DMSO or RP6306 (2μM) for 48h. qPCR was used to detect the mRNA changes of OAS1, CCL2, ISG15, ISG20, IFIT1 (n=3). Two-way ANOVA was used. *P<0.05, **P<0.01, ***P<0.001, ****P<0.0001.

